# The yeast ISW1b ATP-dependent chromatin remodeler is critical for nucleosome spacing and dinucleosome resolution

**DOI:** 10.1101/2020.08.14.251702

**Authors:** Peter R. Eriksson, David J. Clark

## Abstract

Isw1 and Chd1 are ATP-dependent nucleosome-spacing enzymes required to establish regular arrays of phased nucleosomes near transcription start sites of yeast genes. Cells lacking both Isw1 and Chd1 have extremely disrupted chromatin, with weak phasing, irregular spacing and a propensity to form closed-packed dinucleosomes. The Isw1 ATPase subunit occurs in two different remodeling complexes: ISW1a (composed of Isw1 and Ioc3) and ISW1b (composed of Isw1, Ioc2 and Ioc4). The Ioc4 subunit of ISW1b binds preferentially to the H3-K36me3 mark. Here we show that ISW1b is primarily responsible for setting nucleosome spacing and resolving close-packed dinucleosomes, whereas ISW1a plays only a minor role. ISW1b and Chd1 make additive contributions to dinucleosome resolution, such that neither enzyme is capable of resolving all dinucleosomes on its own. Loss of the Set2 H3-K36 methyltransferase partly phenocopies loss of Ioc4, resulting in increased dinucleosome levels with only a weak effect on nucleosome spacing, suggesting that Set2-mediated H3-K36 trimethylation contributes to ISW1b-mediated dinucleosome separation. The H4 tail domain is required for normal nucleosome spacing but not for dinucleosome resolution. We conclude that the nucleosome spacing and dinucleosome resolving activities of ISW1b and Chd1 are critical for normal global chromatin organisation.

## Introduction

Eukaryotic DNA is packaged into the nucleus in the form of chromatin. The structural subunit of chromatin is the nucleosome, which is composed of an octamer of core histones (two molecules each of H3, H4, H2A and H2B), around which is wrapped ^~^146 bp of DNA in ^~^1.7 superhelical turns ^1^. Nucleosomes are regularly spaced along the DNA, like beads on a string, forming a fibre which spontaneously folds into higher-order chromatin structures ^2^. Nucleosomes restrict access to DNA and are potent inhibitors of transcription and other DNA-dependent processes in vitro. Cells regulate access to their DNA in part by deploying ATP-dependent chromatin remodeling complexes that are capable of overcoming the nucleosome, either by removing it from the DNA or by sliding it along the DNA ^3–6^.

The ISWI and CHD enzymes represent a major class of ATP-dependent chromatin remodelers conserved from yeast to mammals. They are primarily nucleosome sliding enzymes; many have nucleosome spacing activity in vitro ^7–13^. In vivo, ISWI enzymes are important for chromatin organization near promoters and other gene regulatory elements ^14–17^. ISWI complexes have additional functions in chromatin assembly ^8,10,18^, stress-induced gene repression ^19,20^, transcript termination ^21,22^ and quality control of mRNP biogenesis ^23^.

The budding yeast (*Saccharomyces cerevisiae*) possesses at least four ATP-dependent chromatin remodeling complexes capable of spacing nucleosomes in vitro: ISW1, ISW2, Chd1 and INO80 ^7,24–26^. In vivo, global chromatin organization in cells lacking Isw2 is very similar to wild type ^14,15^, suggesting that ISW2 activity is more local than global. In contrast, cells lacking Ino80 ^27–29^ or Isw1 ^15,30^ have shorter average nucleosome spacing than wild type cells. Cells lacking Chd1 have slightly shorter spacing and relatively poor nucleosome phasing ^14,15^. Most impressively, cells lacking both Isw1 and Chd1 have extremely disrupted chromatin, indicating that both enzymes are required for proper chromatin organization ^14,15^. Recently, we showed that an important contributory factor to chromatin disruption in the *chd1Δ isw1Δ* double mutant is a tendency for nucleosomes at the 5’-ends of yeast genes to be packed close together, resulting in dinucleosomes with little or no linker DNA ^22^. This observation suggests that Isw1 and/or Chd1 are critical for resolving dinucleosomes.

The yeast Isw1 ATPase subunit is found in two different complexes, termed ISW1a (containing Isw1 and Ioc3) and ISW1b (containing Isw1, Ioc2 and Ioc4) ^26,31^. There is genetic evidence for antagonistic interactions between ISW1a and ISW1b, suggesting that ISW1a has a negative role in transcription that is suppressed by ISW1b ^32^. The Ioc subunits appear to have regulatory functions, since the isolated Isw1 subunit is inactive in vitro ^26^, unless the AutoN inhibitory domain is mutated ^33^. The ISW1a complex is a potent nucleosome spacing enzyme in vitro ^26,34,35^ and its structure has been determined, suggesting a mechanism involving separation of two nucleosomes using a protein ruler ^36^. The ISW1a complex has higher nucleosome spacing activity than the ISW1b complex in vitro ^26^ and contributes much more than ISW1b to nucleosomal array formation at promoters in a purified system ^25^. Isw1 binding to chromatin is indirectly mediated by H3-K4 trimethylation ^37^. The Ioc4 subunit of ISW1b has a PWWP domain which binds to H3-K36me3, a histone modification associated with active transcription ^38,39^. The other auxiliary subunit of ISW1b, Ioc2, contains a putative PHD finger, which may bind to a methylated histone residue ^38^. These observations suggest that H3-K4 and H3-K36 trimethylation may play a critical role in ISW1 function.

Although ISW1a and ISW1b have been studied in depth in vitro, relatively little is known about their contributions to ISW1 function in vivo. Here we have assessed the contributions of ISW1a and ISW1b to nucleosome spacing and separation of dinucleosomes. We find that ISW1b is the major spacing enzyme, whereas ISW1a plays a very minor role in global genic chromatin organization that is revealed only in the absence of ISW1b. ISW1b, together with Chd1, is required to resolve dinucleosomes, whereas ISW1a makes little contribution.

## Results

### The ISW1b complex is the major nucleosome spacing enzyme in yeast

We have used MNase-seq to determine the chromatin organization at yeast genes. Briefly, this technique involves micrococcal nuclease (MNase) digestion of the chromatin in purified nuclei to predominantly mononucleosomes. Suitably digested DNA samples are then used to construct libraries for paired-end sequencing. Alignment to the yeast genome indicates the location of each sequenced nucleosome. The midpoint of each sequence is assumed to represent the midpoint of the nucleosome (also called the dyad). An average plot for all 5770 yeast genes is obtained by aligning all genes on their transcription start site (TSS) or on their major +1 nucleosome position, summing all of the nucleosome dyads on every gene, and normalizing to the genomic average (set at 1). A typical plot for wild type cells is shown in Fig. 1a. We define nucleosome spacing as the average distance between the dyad peaks for the +1 to +5 nucleosomes (i.e. the first five nucleosomes on the average gene), measured by the slope of the regression line. However, in some mutants, the phasing is so poor that the spacing cannot be measured accurately (see below). The degree of phasing is indicated by the height of the nucleosome peaks: higher peaks indicate more coincident dyads and therefore better nucleosome positioning.

**Fig. 1.**
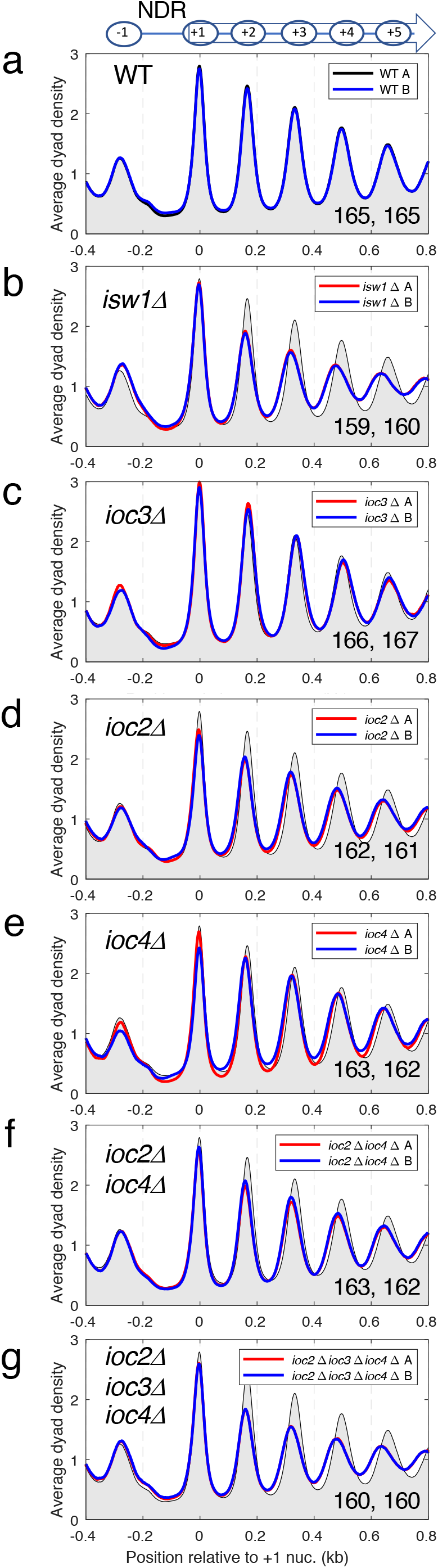
The ISW1b complex (Isw1-Ioc2-Ioc4) is required for setting wild type nucleosome spacing, whereas the ISW1a complex (Isw1-Ioc3) sets intermediate spacing in the absence of ISW1b. Average nucleosome dyad density plots for all genes: (**a**) wild type, (**b**) *isw1Δ*, (**c**) *ioc3Δ*, (**d**) *ioc2Δ*, (**e**) *ioc4Δ*, (**f**) *ioc2Δ ioc4Δ*, (**g**) *ioc2Δ ioc3Δ ioc4Δ*. All yeast genes were aligned on the midpoints of their +1 nucleosomes. The dyad distribution was normalized to the global average (set at 1). For ease of comparison, wild type (WT) replicate A is shown as a black line with grey fill in all plots. Two biological replicate experiments (A and B) are shown for each strain in all of the plots (A: red line; B: blue line). The average spacing in bp is shown for replicates A and B in the bottom right corner (measured by regression analysis of the first 5 nucleosome peaks, beginning with the +1 nucleosome).

We have shown previously ^15^ that cells lacking Isw1 have weaker nucleosome phasing and reduced spacing relative to wild type cells (Fig. 1b). However, *isw1Δ* cells lack both the ISW1a and the ISW1b remodeling complexes. To assess the relative contributions of ISW1a and ISW1b to phasing and spacing, we examined the chromatin organization in various *ioc* mutants. Cells lacking the ISW1a complex (*ioc3Δ*) show a slight increase in spacing (replicates: 166, 167 bp) with no change in phasing relative to wild type (165, 165 bp) (Fig. 1c). In contrast, cells lacking an ISW1b subunit (*ioc2Δ* or *ioc4Δ*) have intermediate nucleosome spacing (161-163 bp) (Fig. 1d, e): less than in wild type (165 bp), but not as low as in isw1Δ cells (159, 160 bp). The spacing differences are mostly small, but they are clear, because the nucleosome shift increases with the nucleosome number (compare biological replicate experiments; Fig. 1). Phasing is weaker in the *ioc2Δ* mutant (Fig. 1d), but hardly affected in the *ioc4Δ* mutant (Fig. 1e). Cells lacking both ISW1b ancillary subunits (the *ioc2Δ ioc4Δ* double mutant) have similar chromatin organization to the *ioc2Δ* single mutant, exhibiting the weaker phasing observed in *ioc2Δ* cells as well as the shorter spacing observed in both the *ioc2Δ* and *ioc4Δ* single mutants (Fig. 1f). The similar spacing in the *ioc2Δ* and *ioc4Δ* single mutants and the *ioc2Δ ioc4Δ* double mutant (^~^162 bp) suggests that both Ioc2 and Ioc4 are required for ISW1b spacing activity.

Chromatin organisation in cells lacking all three ancillary subunits (the *ioc2Δ ioc3Δ ioc4Δ* triple mutant) is very similar to that in *isw1Δ* cells (Fig. 1g). This is expected, because both complexes should be inactive in both mutants, given that the Isw1 subunit by itself has no remodeling activity in vitro ^26^, but it also implies that ISW1a affects spacing since the triple mutant is not the same as the *ioc2Δ ioc4Δ* double mutant. The shorter spacing in the *ioc2Δ ioc3Δ ioc4Δ* triple mutant suggests that ISW1a is responsible for the intermediate spacing observed in the absence of ISW1b. Since the loss of Ioc3 has only a small effect on global spacing in the presence of ISW1b (the *ioc3Δ* single mutant), ISW1b is more important for chromatin organization than ISW1a. That is, the contribution of ISW1a is only clear at the global level if ISW1b is absent. We conclude that the ISW1b complex is primarily responsible for wild type nucleosome spacing and that the ISW1a complex has a minor underlying role, mediating intermediate spacing.

### ISW1a and ISW1b spacing activities are not restricted to genes enriched in their respective Ioc subunits

Chromatin immunoprecipitation (ChIP-seq) experiments have shown that ISW1a (Ioc3), ISW1b (Ioc4) and Isw1 are enriched on different sets of genes, suggesting that these genes might be differentially affected by ISW1 or ISW1b ^27^. We determined whether the chromatin organization of these sets of genes is differentially affected by loss of ISW1a or ISW1b, as might be expected. In the case of ISW1a, we found that Ioc3-enriched genes show very slightly increased spacing (166 vs. 165 bp) in wild type cells and in *ioc3Δ* cells (167 vs. 166 bp), but these differences are probably negligible (Supplementary Fig. S1). Thus, ISW1a has little or no differential effect on the chromatin organization of its putative target genes. Moreover, the spacing on Ioc3-bound genes is affected in *ioc4Δ* cells, indicating that ISW1a-enriched genes are affected by ISW1b (Ioc4) (Supplementary Fig. S1). In the case of ISW1b, Ioc4-enriched and non-enriched genes have the same spacing and phasing in wild type cells, suggesting that they are not differentially affected by ISW1b. In *ioc4Δ* cells, both sets of genes have shorter spacing than wild type (Supplementary Fig. S1). Ioc4-enriched genes may have slightly shorter spacing than non-enriched genes, but the effect is subtle (Supplementary Fig. S1). As expected, Ioc4-enriched genes are not affected in *ioc3Δ* cells (Supplementary Fig. S1). Thus, ISW1b affects the chromatin organization of both sets of genes. Finally, Isw1-enriched genes have the same spacing as non-enriched genes in wild type cells and, although the phasing is slightly better on the Isw1-enriched genes, this effect is also observed in the absence of Isw1 (Supplementary Fig. S1). Isw1-enriched and non-enriched genes show similar changes in *isw1Δ* cells (short spacing and weaker phasing), although there is a slight difference in spacing of 1-2 bp between the two sets of genes (Supplementary Fig. S1). Overall, the Isw1 complexes affect the chromatin organization of both enriched and non-enriched genes similarly, with only subtle differences at most. It is unclear why we do not observe obvious differences in chromatin organization for putative target and non-target genes defined by Ioc3, Ioc4 or Isw1 enrichment; perhaps all genes have some bound ISW1b, such that the putative target genes are only modestly enriched relative to non-target genes.

### ISW1b and Chd1 account for the extreme chromatin disruption in cells lacking both Isw1 and Chd1

Cells lacking Chd1 have a mild chromatin organization defect ^14,15^, characterized by somewhat shorter spacing than wild type (164, 163 bp vs. 165, 165 bp) and weaker phasing (Fig. 2a). In contrast, cells lacking both Chd1 and Isw1 (the *chd1Δ isw1Δ* double mutant) have extremely disrupted chromatin organization ^14,15^ (Fig. 2b). We observed that the *chd1Δ ioc2Δ ioc4Δ* triple mutant (Fig. 2c) also has extremely disrupted chromatin, whereas the *chd1Δ ioc3Δ* double mutant is very similar to the *chd1Δ* single mutant (Fig. 2d). This observation is consistent with our conclusion that the ISW1b complex is much more important than the ISW1a complex for global nucleosome spacing (Fig. 1). However, chromatin organization in the *chd1Δ ioc2Δ ioc4Δ* triple mutant is not quite as severely disrupted as in the *chd1Δ isw1Δ* double mutant: the phasing in the triple mutant is very poor, but somewhat stronger than in the double mutant (Fig. 2b, c). Measurement of the spacing in these two mutants is not appropriate because most of the peaks are too weak to measure accurately. The difference between the *chd1Δ ioc2Δ ioc4Δ* triple mutant and the *chd1Δ isw1Δ* double mutant is that the ISW1a complex (Isw1-Ioc3) is still present in the former mutant. This suggests that ISW1a contributes some residual order to genic chromatin in the absence of both Chd1 and ISW1b.

**Fig. 2.**
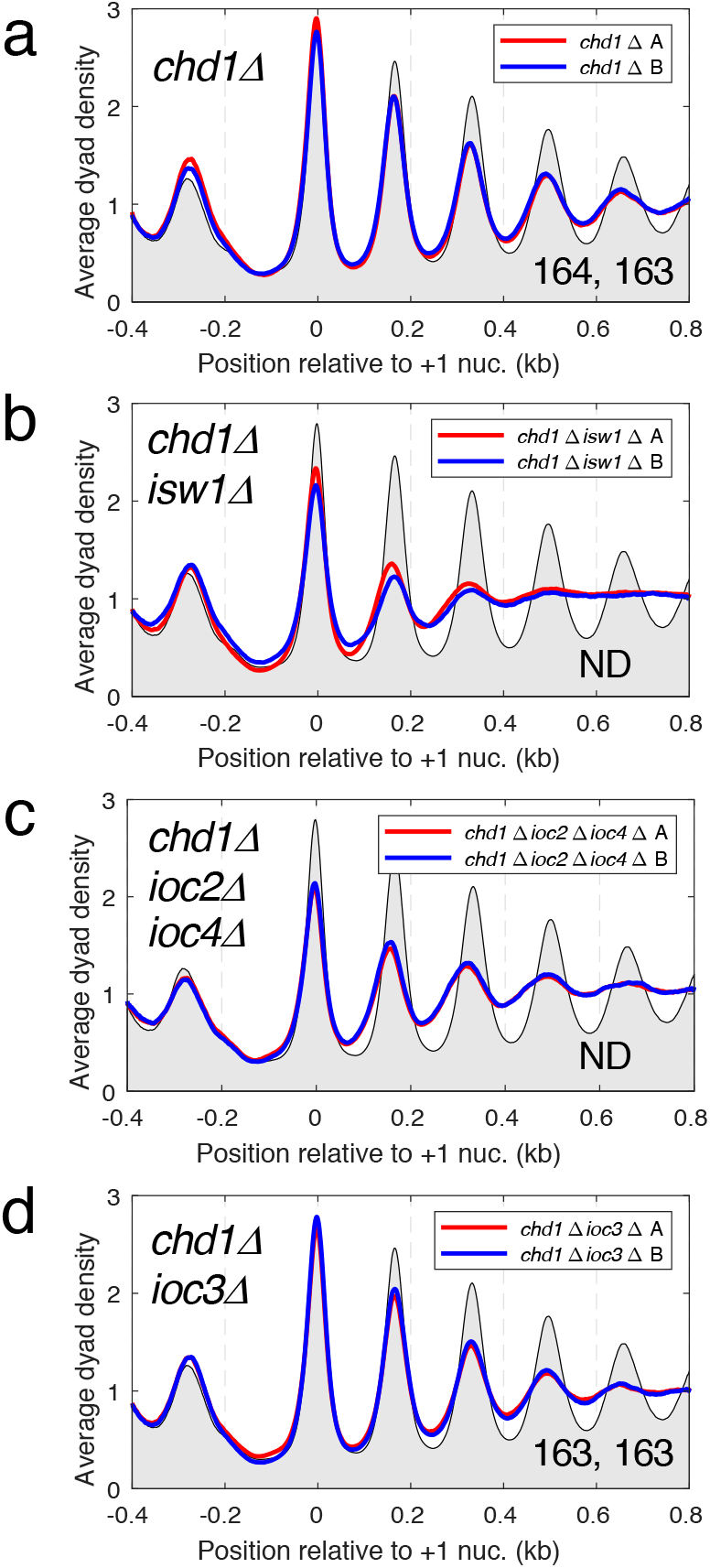
ISW1b and Chd1 are the major remodelers required for normal chromatin organization. Average nucleosome dyad density plots for all genes: (**a**) *chd1Δ*, (**b**) *chd1Δ isw1Δ*, (**c**) *chd1Δ ioc2Δ ioc4Δ*, (**d**) *chd1Δ ioc3Δ*. All yeast genes were aligned on the midpoints of their +1 nucleosomes. The dyad distribution was normalized to the global average (set at 1). For ease of comparison, wild type replicate A is shown as a black line with grey fill in all plots. Two biological replicate experiments (A and B) are shown for each strain in all of the plots (A: red line; B: blue line). The average spacing in bp is shown for replicates A and B in the bottom right corner (measured by regression analysis of the first 5 nucleosome peaks, beginning with the +1 nucleosome); ND = not determined because the phasing is too weak for accurate measurement.

### ISW1a and ISW1b have little effect on the chromatin of very active genes

Since ISW1b preferentially associates with transcriptionally active genes ^39^, we examined whether ISW1a or ISW1b mediate specific effects at highly active genes. We defined highly active genes as those which have > 4 times the genomic average signal using our published ChIP-seq data for the Rpb3 subunit of Pol II ^15^. Although this is a somewhat arbitrary threshold, it is clear from heat map analysis that relatively few genes have high levels of Pol II; most genes have relatively low Pol II levels (Supplementary Fig. S2). On average, the 300 most active genes have ^~^8-fold higher Rpb3 density than the remaining genes. Accordingly, we compared the average chromatin structure of the most active genes with that of the remaining,much less active, 5470 genes (Supplementary Fig. S2). In wild type cells, the chromatin of the highly active genes is poorly organized, characterized by much reduced and irregular spacing, weak phasing and a much wider nucleosome-depleted region (NDR) at the promoter, which extends upstream ^40,41^. Although the average spacing on the highly active genes cannot be measured accurately due to poor phasing, it is clearly shorter than the spacing on the other genes (Supplementary Fig. S2; compare nucleosome peak locations). The chromatin organization of the highly active genes is not strongly differentially affected by any of the *iocΔ* mutations or *isw1Δ* (Supplementary Fig. S2), whereas the less active genes have the spacing observed for all genes, as expected. We conclude that ISW1a and ISW1b have no differential effect on the chromatin organization of highly active genes, with the caveat that their general state of disruption might obscure subtle effects. On the other hand, nucleosome phasing on the top 300 active genes is weaker in cells lacking Chd1 than in wild type or in any of the *ioc* mutants (Supplementary Fig. S2), suggesting that Chd1 is the most important spacing enzyme for highly active genes, and consistent with a direct association of Chd1 with transcript elongation factors ^42^.

### Both ISW1b and Chd1 are important for resolution of close-packed dinucleosomes

We reported previously that a major contributing factor to chromatin disruption in the *chd1Δ isw1Δ* double mutant is the presence of close-packed dinucleosomes, primarily involving the +2 nucleosome (i.e. dinucleosomes containing either the +1 and +2 nucleosomes or the +2 and +3 nucleosomes) ^22^ (Fig. 3a). These dinucleosomes are characterized by MNase-resistant DNA fragments of 250 - 350 bp, presumably representing two nucleosomes (or perhaps sub-nucleosomes) with little or no intervening linker DNA for MNase to cut. The dyad plots shown above (Figs. 1 and 2) include only mononucleosome data (the analysis was limited to DNA fragments of 120 - 180 bp). A dyad plot is not appropriate for dinucleosomes, since the midpoint of a dinucleosome would be located between the two nucleosomes. Instead, we used nucleosome occupancy (coverage) plots, in which the number of times each genomic base pair appears in either mononucleosomes or dinucleosomes is counted and normalized to the genomic average.

**Fig. 3.**
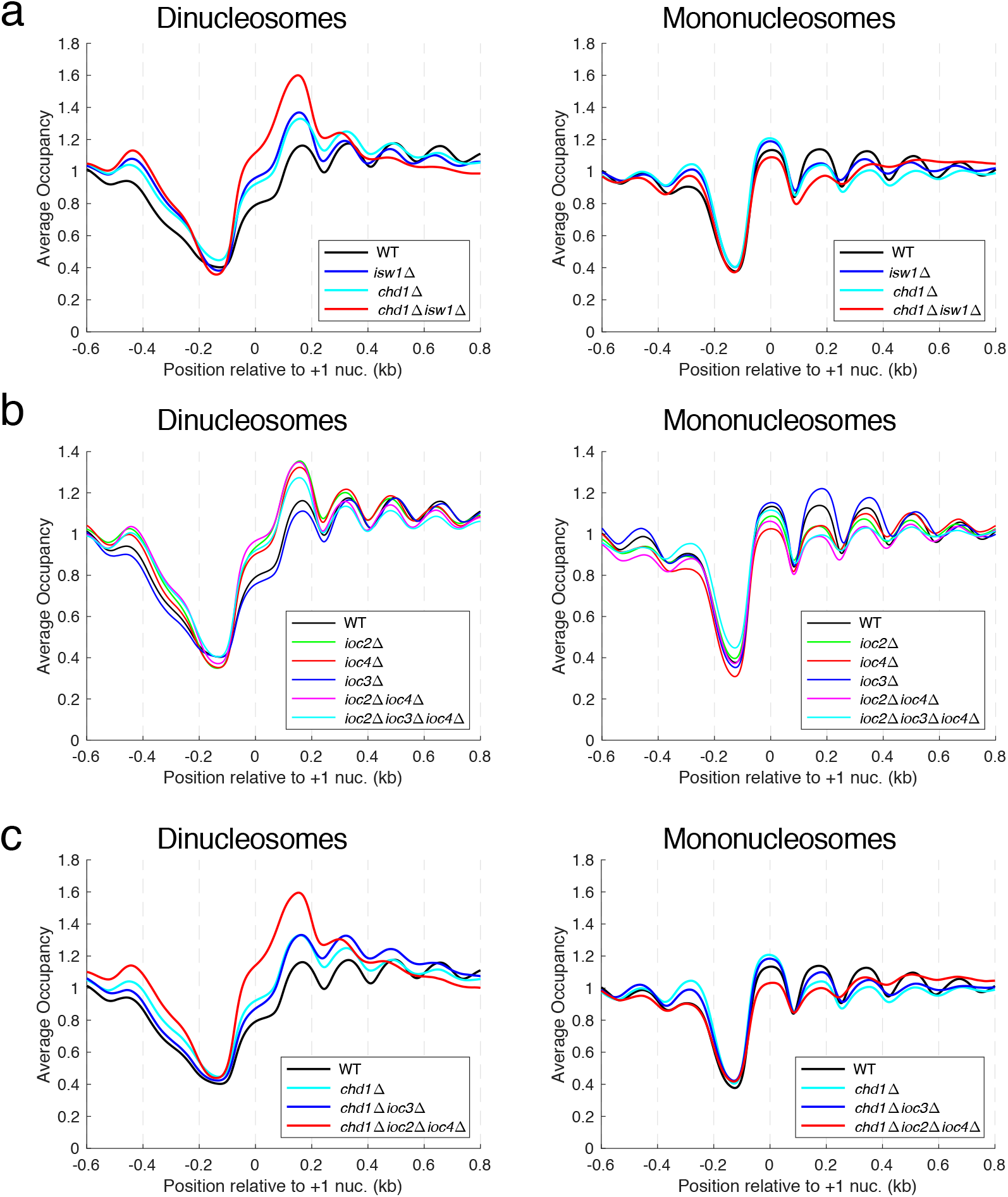
Both ISW1b and Chd1 are important for separating close-packed dinucleosomes. Average nucleosome occupancy (coverage) plots for all genes. All yeast genes were aligned on the midpoint of their average +1 nucleosome position. The occupancy was normalized to the global average (set at 1) for mononucleosomes (120 - 180 bp) or dinucleosomes (250 - 350 bp). Note different y-axis scales are used in different plots to separate the lines more clearly. Data for replicate A are shown (see Supplementary Fig. S3 for comparison of biological replicate experiments). **(a)** Dinucleosomes involving the +2 nucleosome are enriched whereas +2 mononucleosomes are depleted in *chd1Δ isw1Δ* cells; smaller effects are observed in *chd1Δ* and *isw1Δ* cells. **(b)** ISW1b (Isw1-Ioc2-Ioc4) accounts for the increased level of dinucleosomes in *isw1Δ* cells. **(c)** ISW1b and Chd1 are both required to resolve all dinucleosomes.

In the *chd1Δ isw1Δ* double mutant, there is a strong dinucleosome peak at the +2 position and a depressed +2 mononucleosome peak, consistent with the presence of a significant fraction of all +2 nucleosomes in dinucleosomes ^22^ (Fig. 3a; Supplementary Fig. S3). We determined the relative contributions of Chd1 and Isw1 to dinucleosome resolution. A dinucleosome peak is observed in both the *chd1Δ* and *isw1Δ* single mutants, but it is weaker than in the *chd1Δ isw1Δ* double mutant (Fig. 3a). The +2 mononucleosome peak is also weaker in the single mutants relative to wild type. These data indicate that the Chd1 and Isw1 remodeling enzymes both contribute independently and make additive contributions to dinucleosome resolution, such that neither enzyme is capable of resolving all dinucleosomes on its own.

Next, we determined the separate contributions of the ISW1a and ISW1b complexes to Isw1-dependent dinucleosome resolution (Fig. 3b). The *ioc2Δ* and *ioc4Δ* single mutants have an enhanced dinucleosome peak and depressed mononucleosome peak, as observed for the isw1Δ single mutant, suggesting that both ancillary subunits of the ISW1b complex are required for Isw1-dependent dinucleosome resolution. Consistent with this conclusion, the *ioc2Δ ioc4Δ* double mutant has a dinucleosome peak similar to that in the single mutants. In contrast, there are fewer dinucleosomes in the *ioc3Δ* mutant, which has a slightly lower +2 dinucleosome peak than wild type and somewhat higher levels of the +2 and +3 mononucleosomes (Fig. 3b; Supplementary Fig. S3), indicating that ISW1a is not important for dinucleosome resolution. These conclusions are supported by high levels of dinucleosomes in the *chd1Δ ioc2Δ ioc4Δ* triple mutant versus wild type, resulting from loss of both ISW1b and Chd1 (Fig. 3c). Similarly, the *chd1Δ ioc3Δ* double mutant has more dinucleosomes than wild type, but less than the *chd1Δ ioc2Δ ioc4Δ* triple mutant, which can be accounted for by the absence of Chd1 with little contribution from ISW1a (Fig. 3c). Thus, the ISW1b complex accounts quite well for the role of Isw1 in dinucleosome resolution.

### Set2 contributes to dinucleosome separation

The Ioc4 PWWP domain binds preferentially to H3-K36me3 relative to unmethylated H3-K36 in vitro, suggesting that ISW1b may be regulated by H3-K36me3 ^38,39^. The only H3-K36 methyltransferase in yeast is encoded by *SET2*. We examined whether ISW1b-mediated changes in nucleosome spacing and dinucleosome resolution depend on Set2. Nucleosome spacing in a *set2Δ* mutant (replicates: 165 and 163 bp) is slightly lower than wild type (165 and 165 bp), although this difference is probably not significant (Fig. 4a). Loss of Ioc4 has a stronger effect on nucleosome spacing (Fig. 1e; *ioc4Δ* replicates: 163 and 162 bp) than loss of Set2 (Fig. 4b). Like *ioc4Δ* cells, *set2Δ* cells have higher dinucleosome levels than wild type cells (Fig. 4c). Interestingly, there are more +3 and +4 dinucleosomes in *set2Δ* cells than in *ioc4Δ* cells (Fig. 4c; Fig. 3b) Supplementary Fig. S3). However, there is no corresponding depression in the +2, +3 and +4 mononucleosome peaks in *set2Δ* cells (Fig. 4c); instead, these mononucleosomes are higher than in wild type. There is an apparently compensatory reduction in NDR occupancy and the −1 nucleosome (Fig. 4c), which is also observed in *ioc4Δ* cells but not in the other *ioc* mutants (Fig. 3b). Overall, these data suggest that Set2 and H3-K36me3 may be more important for the dinucleosome resolving activity of ISW1b than for its spacing activity.

**Fig. 4.**
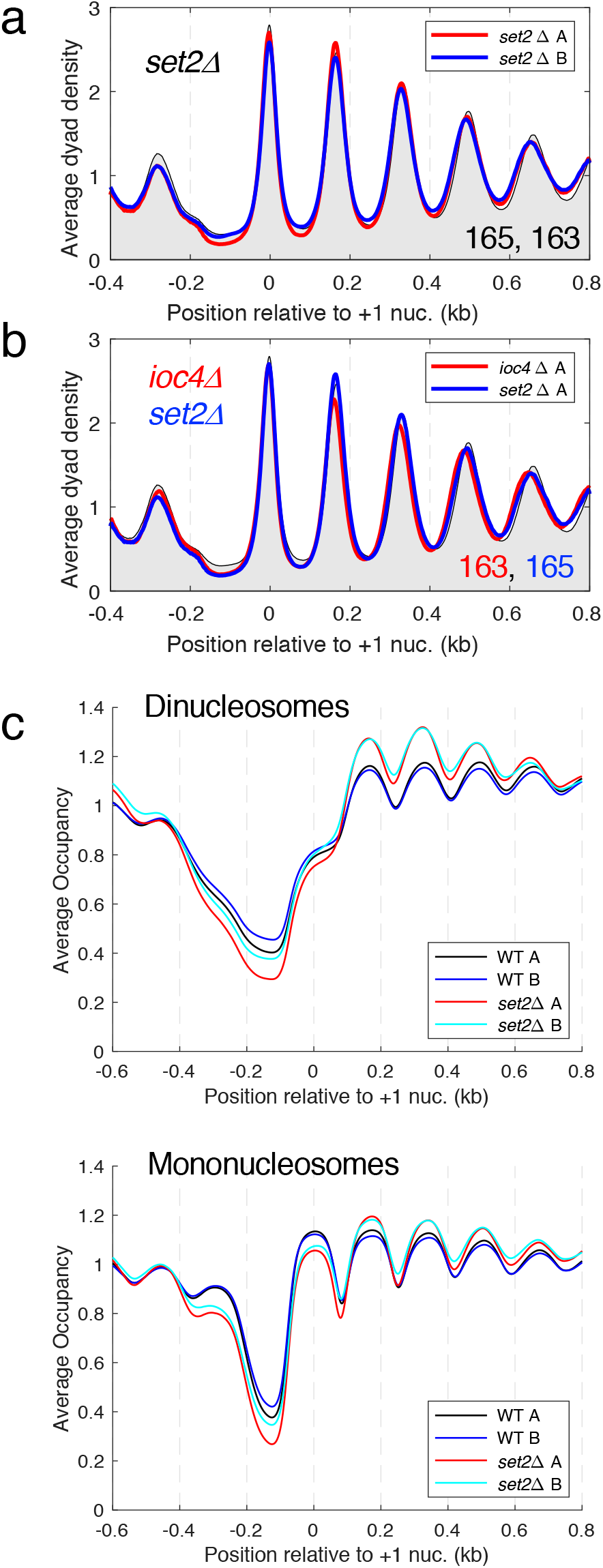
Set2 contributes to dinucleosome separation but has little effect on spacing. (**a**) Average nucleosome dyad density plot for all genes in *set2Δ* cells. Wild type replicate A is shown as a black line with grey fill. Two biological replicate experiments are shown: A (red line) B (blue line). The average spacing (bp) for replicates A and B is shown (bottom right). (**b**) Comparison of chromatin organization in *set2Δ* cells (blue line) and *ioc4Δ* cells (red line) (replicates A). (**c**) Occupancy plots for dinucleosomes and mononucleosomes in *set2Δ* and wild type (WT) cells (see legend to Fig. 3).

### Loss of Set1 has no effect on global chromatin organisation

The binding of ISW1 to chromatin is mediated indirectly through H3-K4 trimethylation, ^37^. On the other hand, a direct interaction might be possible, mediated by the putative PHD domain in the Ioc2 subunit of ISW1b ^38^. Since Set1 is the only H3-K4 methyltransferase in yeast, we tested this possibility by examining *set1Δ* cells. However, chromatin organization in *set1Δ* cells is essentially identical to wild type at the global level (Supplementary Fig. S4) and quite different from that in *ioc2Δ* cells (Fig. 1d). Dinucleosomes involving the +3 and +4 nucleosomes may be somewhat elevated in *set1Δ* cells, although this is unclear because the replicates are not consistent in this respect (Supplementary Fig. S3; Supplementary Fig. S4). In conclusion, Set1 and therefore H3-K4me3 are not required for ISW1-dependent nucleosome spacing.

### Deletion of the H4 N-terminal tail domain results in shorter nucleosome spacing but does not increase dinucleosome levels

Nucleosome mobilisation in vitro by ISWI complexes isolated from different organisms requires the H4 N-terminal tail domain, specifically the “basic patch” (residues 16 to 19) ^33,43,44^. In addition, genetic interactions between *isw1* mutations and H4 mutations (point mutations and a tail deletion) suggest involvement in a common pathway ^19^. We reasoned that deletion of the H4 N-tail might result in a chromatin organization similar to that observed in the *isw1Δ* single mutant: reduced spacing, weaker phasing and increased dinucleosome formation (Fig 1b; Fig. 3a). We constructed a yeast strain in which both H3/H4 gene loci (*HHT1-HHF1* and *HHT2-HHF2*) were deleted and covered by a plasmid carrying wild type *HHT1-HHF1* or *HHT1-HHF1Δ21*, in which the first 21 amino acid residues of the H4 N-tail had been deleted. This strain displayed a clear growth phenotype, with a doubling time of ^~^2.6 h, compared with ^~^1.8 h for the wild type strain, in synthetic complete (SC) medium. We find that nucleosome spacing is shorter in cells lacking the H4 N-tail (Fig. 5a), though not quite as short as in *isw1Δ* cells (Fig. 1b), and both mutants have poor phasing. These observations are consistent with the requirement of Isw1 for the H4 N-tail domain. However, removal of the H4 N-tail resulted in only a very slight increase in dinucleosome prevalence (Fig. 5b; Supplementary Fig. S3), unlike loss of Isw1 (Fig. 3a), indicating that loss of Isw1 and loss of the H4 N-tail are not equivalent.

**Fig. 5.**
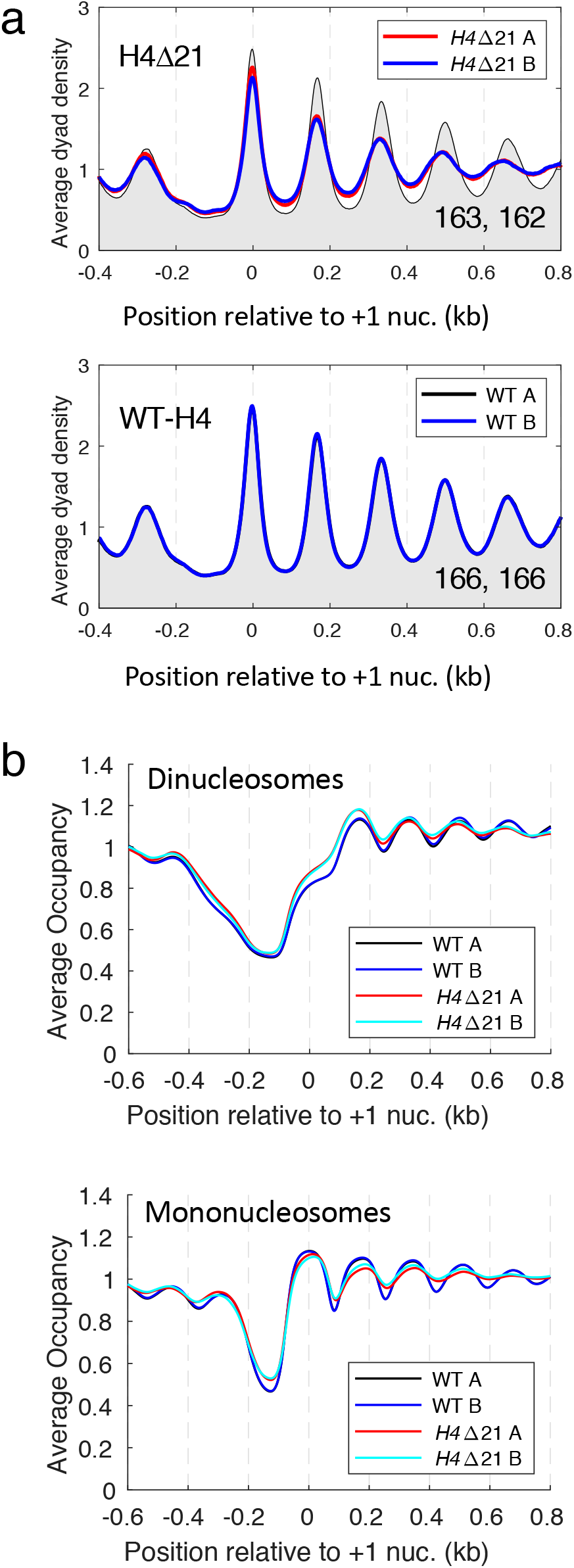
Deletion of the first 21 residues of the H4 N-terminal tail domain results in shorter global nucleosome spacing but little dinucleosome accumulation. **(a)** Comparison of the H4ΔN21 mutant with its isogenic wild type strain (WT-H4) (top panel). WT-H4 replicates (bottom panel). All yeast genes were aligned on the midpoints of their +1 nucleosomes. The dyad distribution was normalised to the global average (set at 1). WT-H4 replicate A: black line with grey fill in both plots. The average spacing in bp is shown for each replicate in the bottom right corner. **(b)** Occupancy plots for dinucleosomes and mononucleosomes in H4Δ21 and wild type (WT) cells (see legend to Fig. 3).

## Discussion

### ISW1b is the primary nucleosome spacing enzyme in yeast

We and others have shown previously that Isw1 and Chd1 are both needed for normal global chromatin organization in yeast ^14,15^. We proposed that these two enzymes compete to set nucleosome spacing in wild type cells, with Isw1 being dominant, setting wild type spacing, and Chd1 directing shorter spacing ^15^. In the absence of Isw1, the spacing is short, which we attributed to Chd1 activity. An important complication, which we have addressed here, is that there are two complexes containing the Isw1 ATPase subunit ^26^. We have determined their respective roles in chromatin organization in vivo: ISW1b and Chd1 are primarily responsible for global nucleosome spacing in wild type cells, whereas a role for ISW1a in directing intermediate spacing only becomes apparent when ISW1b is absent (Fig. 6a). We note that Ino80C also affects global spacing ^27–29^, but it is not yet clear how its activity meshes with those of ISW1b and Chd1.

**Fig. 6.**
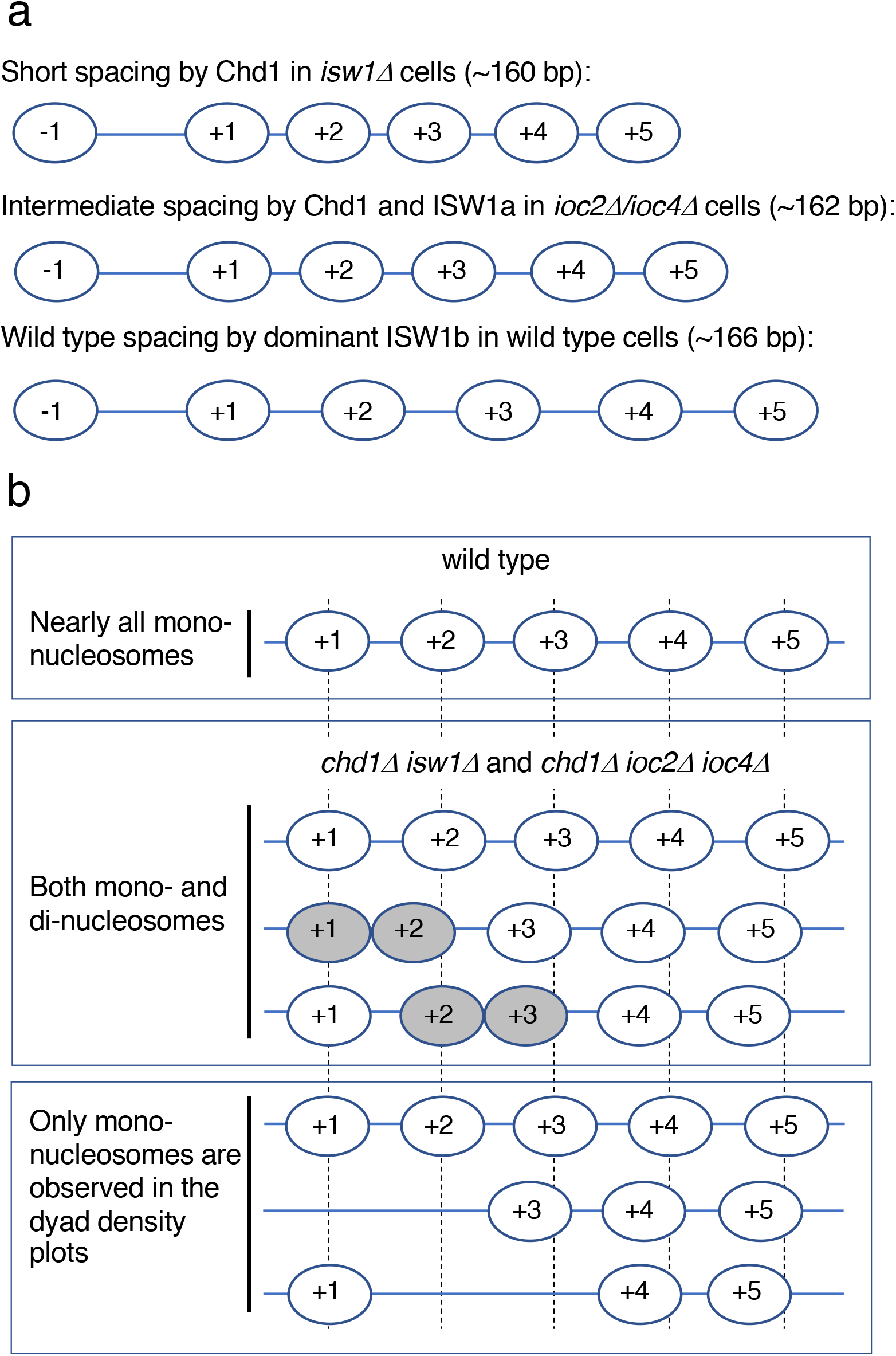
Dominant role of ISW1b in chromatin organization. (**a**) Global average nucleosome spacing is primarily determined by ISW1b (wild type cells), since loss of ISW1a has little effect (*ioc3Δ* mutant). In the absence of ISW1b (*ioc2Δ* and *ioc4Δ* mutants), intermediate spacing is observed, attributed to ISW1a, since the spacing is short in the absence of both ISW1a and ISW1b (*isw1Δ* mutant or *ioc2Δ ioc3Δ ioc4Δ* triple mutant). Short spacing is attributed to Chd1, because the *chd1Δ isw1Δ* double mutant has highly disrupted chromatin. (**b**) Dinucleosomes disrupt phasing and reduce spacing. Model to show the effects of dinucleosomes on chromatin organization. Top panel: Average nucleosome positions on a gene in wild type cells with regular spacing. Middle: Close-packed dinucleosomes (grey ovals) are high in the absence of both ISW1b and Chd1. These dinucleosomes preferentially involve the +2 nucleosome i.e., they are mostly +1/+2 or +2/+3 dinucleosomes. Nucleosomes farther down the gene (+3 etc.) may be regularly spaced relative to the dinucleosome, but because a linker is missing in the dinucleosome, downstream nucleosomes are out of phase with other nucleosomal arrays, resulting in poor phasing due to interference patterns from the different arrays. Bottom: Dinucleosomes are absent from the mononucleosome dyad density plots, resulting in depressed, flattened peaks. These effects are strongest in cells lacking both ISW1b and Chd1. Dinucleosomes that cannot be resolved by ISW1b can account for shorter spacing in *chd1Δ* cells.

A difficulty for our competition model is that, in the absence of Chd1, ISW1b is expected to win the competition, resulting in wild type spacing. Instead, we observe that spacing is somewhat shorter in cells lacking Chd1 ^15^ (Fig. 2a). One possible explanation is that the dominance of ISW1b depends on Chd1 and so, in its absence, we see an increased contribution from ISW1a, such that the spacing represents the average of ISW1a and ISW1b activities. If so, cells having only ISW1b (the *chd1Δ ioc3Δ* double mutant) are expected to have wild type spacing. However, the result is intermediate spacing, very similar to the *chd1Δ* single mutant (Fig. 2). Alternatively, ISW1b might only be able to create nucleosome arrays with wild type spacing if Chd1 has already made arrays with short spacing, although this model is inconsistent with in vitro data showing that purified ISW1b can space nucleosomes by itself ^26^.

A more satisfying explanation involves dinucleosomes (Fig. 6b). Consider a single gene. If there are regularly spaced nucleosomes downstream of a close-packed +1/+2 dinucleosome (i.e. no linker), these nucleosomes will be out of phase by one linker length with nucleosomes in cells with no dinucleosome on this gene. Other cells may have a +2/+3 dinucleosome instead, which will alter the positions of regularly spaced downstream nucleosomes to give a different phasing. The result is generally weaker phasing (peak flattening) and a shift to shorter average spacing. Moreover, since the dinucleosomes are not counted in the mononucleosome phasing pattern, there will be missing occupancy around the +2 nucleosome, resulting in more pattern disruption (primarily a depressed +2 nucleosome peak). These effects will increase as the fraction of genes having a dinucleosome increases: the *chd1Δ isw1Δ* double mutant and the *chd1Δ ioc2Δ ioc4Δ* triple mutant both have high dinucleosome levels and extremely poor phasing. The *chd1Δ* single mutant has fewer dinucleosomes, resulting in a smaller shift in the peaks to shorter spacing. These considerations can account for the shift to shorter spacing in the absence of Chd1 and for the major disruption of chromatin in the absence of both Chd1 and ISW1b. Thus, we propose that the spacing is shorter in cells lacking Chd1 because ISW1b cannot resolve all of the dinucleosomes.

Previously, we showed that Isw1 and/or Chd1 is required to resolve dinucleosomes, or to prevent their formation ^22^. Here we show that both enzymes are important for separating dinucleosomes and that, as observed for spacing activity, ISW1b is much more important than ISW1a. In vitro, both ISW1a and ISW1b can move nucleosomes reconstituted on a pair of 601 nucleosome positioning sequences farther apart, even if separated by a linker of only 4 bp ^45^, indicating that both ISW1 complexes probably have close-packed dinucleosome resolving activity.

### The Set2 methyltransferase contributes to dinucleosome separation

We observe that the ancillary subunits of ISW1b, Ioc2 and Ioc4, are both necessary for ISW1b-dependent chromatin organization, although Ioc2 makes a greater contribution, since it affects phasing as well as spacing. We suggest that Ioc2 and Ioc4 cooperate to influence the remodeling activity of the Isw1 subunit. Both may be linked to specific histone modifications: Ioc4 binds to reconstituted nucleosomes carrying H3-K36me3 with higher affinity than to unmethylated nucleosomes, suggesting that Set2-mediated H3-K36me3 might be an important regulator of ISW1b ^38,39,46^. Our data provide some evidence to support this proposal, because although loss of Set2 has only a subtle effect on nucleosome spacing, it does result in increased dinucleosome levels, similar to that observed for loss of Ioc4. We propose that Set2-mediated trimethylation of H3-K36 is important for ISW1b-mediated dinucleosome resolution, through interaction of H3-K36me3 with its Ioc4 subunit.

We note that neither ISW1 complex requires H3-K36me3 to space nucleosomes in vitro, since they are active on nucleosomes containing unmodified recombinant histones. On the other hand, in vivo, H3-K36me3 may have local effects on ISW1b activity, rather than global effects, which we have not detected. These are probably not occurring at the most active genes because their chromatin organization in *set2Δ* cells is similar to wild type (Supplementary Fig. S2). H3-K36me3 recognition may also affect other functions of ISW1b, such as gene repression ^20^ or mRNP quality control ^23^.

The fact that Ioc2 has a putative PHD domain suggests that ISW1b may interact with an additional methylated histone residue, although there is no evidence for this at present ^38^. It is unlikely to be H3-K4me3, because our data indicate that the nucleosome spacing activity of the ISW1b complex is independent of Set1 and therefore of H3-K4me3. H3-K4me3 may also bind to one of the chromodomains of Chd1 ^47^, although this is controversial ^48^. If Chd1 does indeed bind to H3-K4me3, this binding is not critical for its nucleosome spacing function, since we find that loss of Set1 does not result in global chromatin defects, unlike loss of Chd1.

### The H4 N-terminal tail domain is required for normal nucleosome spacing

The H4 tail domain is generally required by ISWI-like enzymes for activity ^4,5,43,49^. The ISWI ATPase subunit has an inhibitory AutoN domain that resembles the H4 tail, which is displaced by the H4 tail when ISWI binds to a nucleosome ^4,5^. The isolated yeast Isw1 subunit is inactive without Ioc subunits unless the AutoN domain is inactivated by mutation ^33^, implying that the Ioc subunits may regulate Isw1 activity through the AutoN domain. We find that the H4 tail deletion mutant exhibits shorter spacing, consistent with inactivation of ISW1b, but there is little increase in dinucleosomes. The most likely explanation for the non-equivalence of the H4Δ21 and *isw1Δ* mutations is that the H4 tail domain is involved in multiple processes, including interactions with other remodelers and proteins. This possibility is supported by the fact that the H4Δ21 mutant has a strong growth defect, unlike the various *ioc* mutants. Since Chd1 also makes contacts with the basic patch in the H4 tail ^50,51^, its remodeling activity may also be compromised, although H4Δ21 chromatin is clearly not as disrupted as *chd1Δ isw1Δ* chromatin. The H4 N-tail is also required by ISW2 in vitro ^52^, although global chromatin organization is unaffected in *isw2Δ* cells ^14,15^. The low prevalence of dinucleosomes in the H4Δ21 mutant suggests that the H4 tail might be required for dinucleosome formation, possibly because the increased negative charge on nucleosomes lacking the H4 tail might limit how closely nucleosomes can approach one another ^53,54^. Deletion of the H4 tail also results in the loss of multiple post-translational modification sites, preventing interaction with many chromatin factors.

### Data Availability

The datasets generated during the current study are available in the NCBI Gene Expression Omnibus (GEO) repository [GSE156224]. [*Currently private. Reviewer link on front page*].

Published data sets used in the analysis of the current study are also available in the GEO repository (MNase-seq data for YJO484: GSE117514 (Ocampo et al., 2019); Rpb3 ChIP-seq data: GSE69400 ^15^).

## Methods

### Plasmid construction

The primers used in this study are listed in Supplementary Table S1. pRS-HHT1-HHF1 (p368) was constructed by insertion of the 1930-bp SacI-PstI fragment containing the *HHT1-HHF1* locus from p367 ^55^ at the same sites in pRS317 (*CEN ARS LYS2*) (ATCC 77157). pRS-HHT1-HHF1Δ21 (p730) was constructed by replacing the 752-bp AfeI-SacI fragment in p368 with a 689-bp AfeI-SacI fragment with the H4 N-terminal tail deletion (H4 begins Met-Leu22…) made by ligating two PCR fragments together (obtained using p368 as template with primers 1770/1771 and 1773/1812). The sequence was confirmed.

### Yeast strain construction

The yeast strains used in this study are listed in Supplementary Table S2. YDC507 (H4 N-terminal tail deletion) was constructed by transforming ROY1281 (Dhillon and Kamakaka, 2000), which carries plasmid pCC67 (2 micron origin *URA3 HHT1-HHT1 HTA1-HTB1*) (Clark-Adams et al., 1988), with p730 (*CEN ARS LYS2 HHT1-HHF1Δ21*), selection on plates made with synthetic complete medium without lysine (SC-lys) and then counter-selection against *URA3* with 5-fluoro-orotic acid (5-FOA) to evict pCC67. A wild type control strain (YDC101) was constructed in the same way using p368 (*CEN ARS LYS2 HHT1-HHF1*) instead of p730. The diploid strain YPE600 was made by crossing YTT196 ^7^ with YDC111 ^56^. YPE606 (*ioc3Δ*) was constructed by transforming YPE600 with an *ioc3Δ::KanMX* fragment made using primers 1918/1919 and genomic DNA from YTT645, followed by G418 selection on YPD plates and sporulation. YPE608 (*ioc4Δ*) was constructed by transforming YPE600 with an *ioc4Δ::HPH1* fragment made using primers 1922/1923 and genomic DNA from YTT827, followed by hygromycin selection on YPD plates and sporulation. YPE636 (*ioc2Δ*) was constructed by transforming YPE600 with an *ioc2Δ::URA3* fragment made using primers 1961/1962 and extended using primers 1965/1966 and wild type genomic DNA as template, followed by selection on SC-ura plates and sporulation. YPE654 (*ioc2Δ ioc3Δ*) was constructed by transforming YPE636 with the *ioc3Δ::KanMX* DNA fragment (see above). YPE655 (*ioc2Δ ioc4Δ*) was constructed by transforming YPE636 with the *ioc4Δ::HPH1* fragment (see above). YPE657 (*ioc2Δ ioc3Δ ioc4Δ*) was constructed by transforming YPE654 with the *ioc4Δ::HPH1* fragment. YPE712 (*chd1Δ ioc3Δ*) was obtained by crossing YJO486 ^15^ with YPE607. YJO487 was obtained by sporulation of YJO505 ^15^. YPE715 (*chd1Δ ioc2Δ ioc4Δ*) was constructed by crossing YJO487 with YPE655. YPE602 (*set1Δ*) was constructed by transforming YPE600 with a *set1Δ::NAT1* fragment made using primers 1926/1927 and genomic DNA from YTT1986, followed by nourseothricin selection on YPD plates and sporulation. YPE604 (*set2Δ*) was constructed by transforming YDC111 with a *set2Δ::TRP1* fragment made using primers 1914/1915 extended with 1916/17 and plasmid pFA6a-TRP1 ^57^, followed by selection on SC-trp plates.

### MNase-seq

Nuclei were prepared as described ^58^. MNase digestion of nuclei and construction of paired-end libraries was as described ^59^, except that mononucleosomal DNA was not gel-purified, in order to retain mononucleosomes and dinucleosomes in the correct proportions ^22^. Two biological replicate experiments were performed for all strains (i.e., the replicate experiments for each strain were performed entirely independently). MNase-seq data were analysed using scripts originally described by ^22^; modified code is provided (Supplementary Code).

## Acknowledgements

We thank Alan Hinnebusch for helpful comments on the manuscript and Răzvan Chereji for help with the bioinformatics. We thank Alex Naim for constructing p368, Toshi Tsukiyama for yeast strains, and Rohinton Kamakaka and Namrita Dhillon for pCC67 (pRO149) and ROY1281. This research was supported by the Intramural Research Program of the NIH (NICHD). We thank the NHLBI Core Facility (Yan Luo, Poching Liu, and Jun Zhu) for paired-end sequencing. This study utilized the high performance computational capabilities of the Biowulf Linux cluster at the National Institutes of Health.

## Author contributions

PRE: Acquisition, analysis and interpretation of data. DJC: Analysis, interpretation of data and manuscript preparation.

---

**Supplementary Fig. S1.**
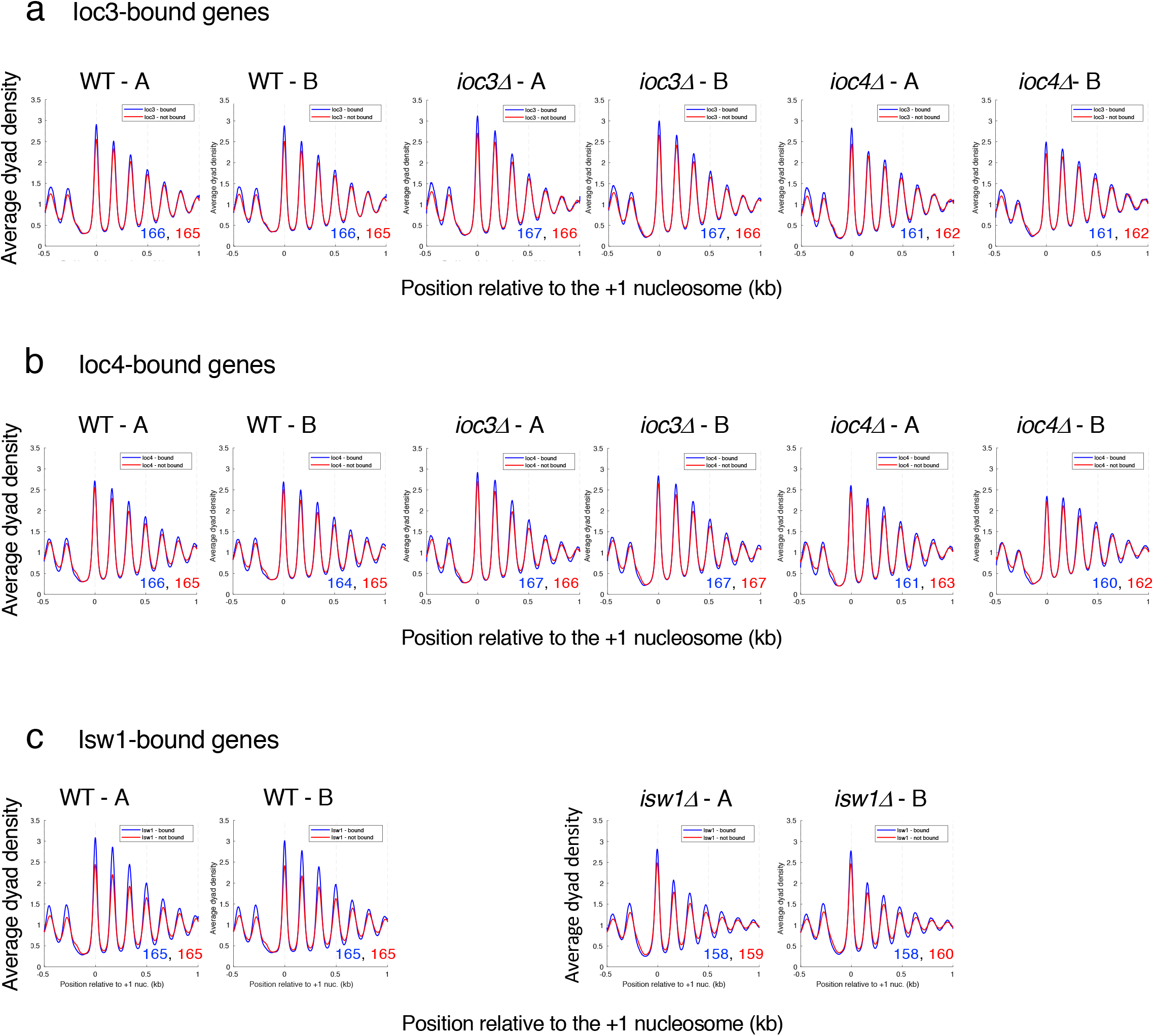
The chromatin organisations of loc3-bound, loc4-bound and Isw1-bound genes are not significantly different from the unbound genes. Genes enriched for ISW1a (Ioc3-bound), ISW1b (Ioc4-bound) and both complexes (Isw1-bound) are defined by ChIP-seq data (Yen et al. 2012). Average nucleosome dyad density plots for bound and unbound genes: (**a**) Ioc3-bound (blue line) and non-bound (red line) genes in wild type (WT) and ioc3Δ cells. (**b**) Ioc4-bound (blue line) and non-bound (red line) genes in WT and ioc4Δ cells. **(c**) Isw1-bound (blue line) and non-bound (red line) genes in WT and isw1Δ cells. All 5770 yeast genes were aligned on the midpoints of their +1 nucleosomes. The dyad distribution was normalised to the global average (set at 1). Two biological replicate experiments (A and B) are shown. The average spacing in bp is shown for bound genes (blue text) and non-bound genes (red text) in the bottom right corner (measured by regression analysis of the first 5 nucleosome peaks, beginning with the +1 nucleosome).

**Supplementary Fig. S2.**
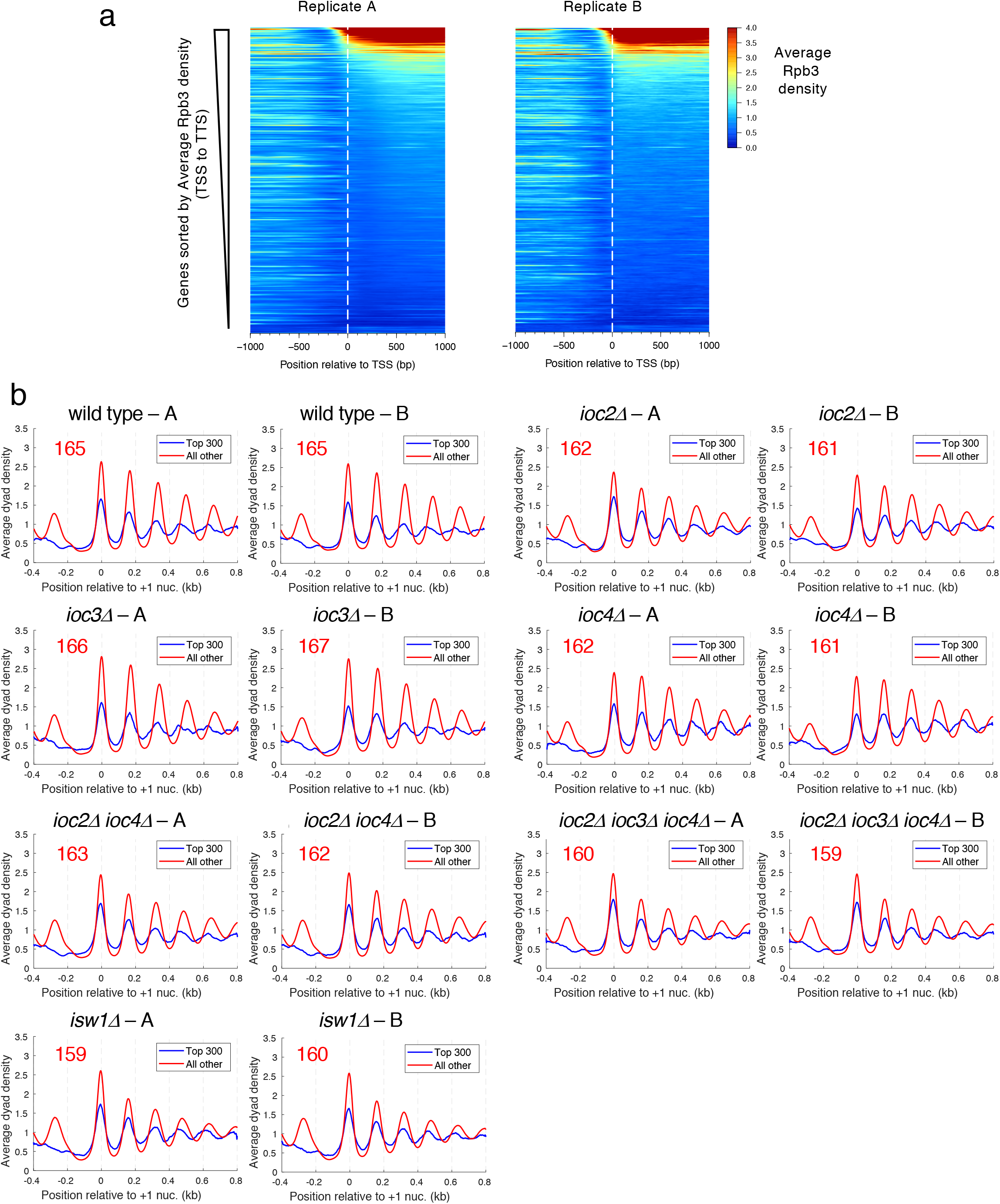
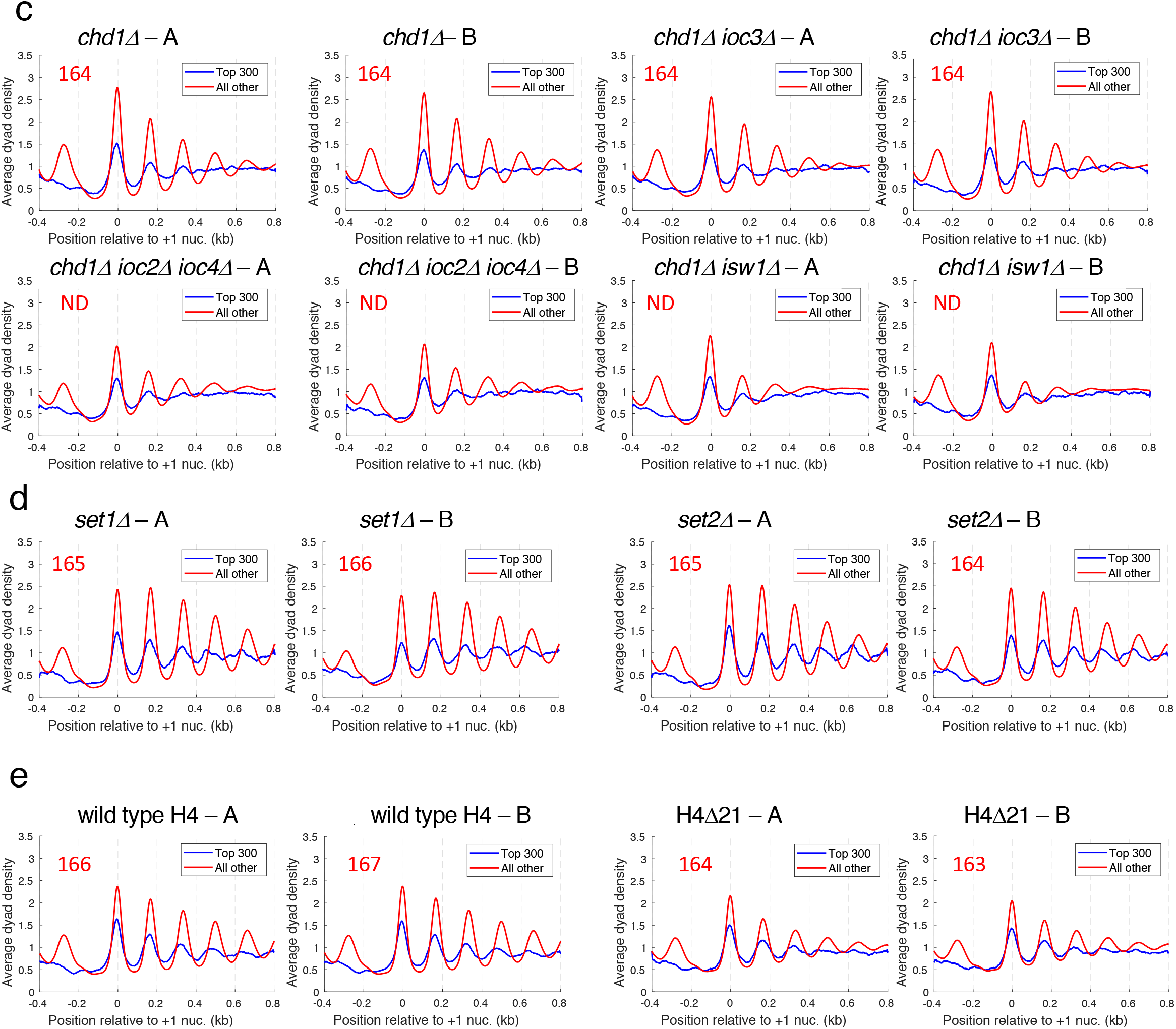
Chromatin organisation of the most highly transcribed genes compared to the remaining genes. (**a**) Heat map analysis of genic Pol II density. ChIP-seq data for the Rpb3 subunit of Pol II in wild type cells (biological replicate data from (Ocampo et al. 2016). The average Pol II density for each of the 5770 genes was computed from transcription start site (TSS) to transcript termination site (TTS). Genes were aligned on the TSS and sorted from high to low Rpb3 density. (**b-e**) Average nucleosome dyad density plots for the 304 most active genes (defined by Rpb3 density > 4 times the genomic average: blue line) compared with the remaining 5470 genes (red line). Genes were aligned on the midpoints of their +1 nucleosomes. The dyad distribution was normalised to the global average (set at 1). Two biological replicate experiments (A and B) are shown in separate panels. The average spacing in bp is shown for the top 300 active genes (blue text) and the other genes (red text), as measured by regression analysis of the first 5 nucleosome peaks, beginning with the +1 nucleosome. **(b**) Wild type (WT), *ioc2Δ, ioc3Δ ioc4Δ, ioc2Δ, ioc2Δ ioc4Δ, ioc2Δ ioc3Δ ioc4Δ* and *isw1Δ*. (**c**) *chd1Δ, chd1Δ ioc3Δ, chd1Δ ioc2Δ ioc4Δ* and *chd1Δ isw1Δ*. (**d**) *set1Δ* and *set2Δ*. (**e**) Wild type H4 and H4Δ21.

**Supplementary Fig. S3.**
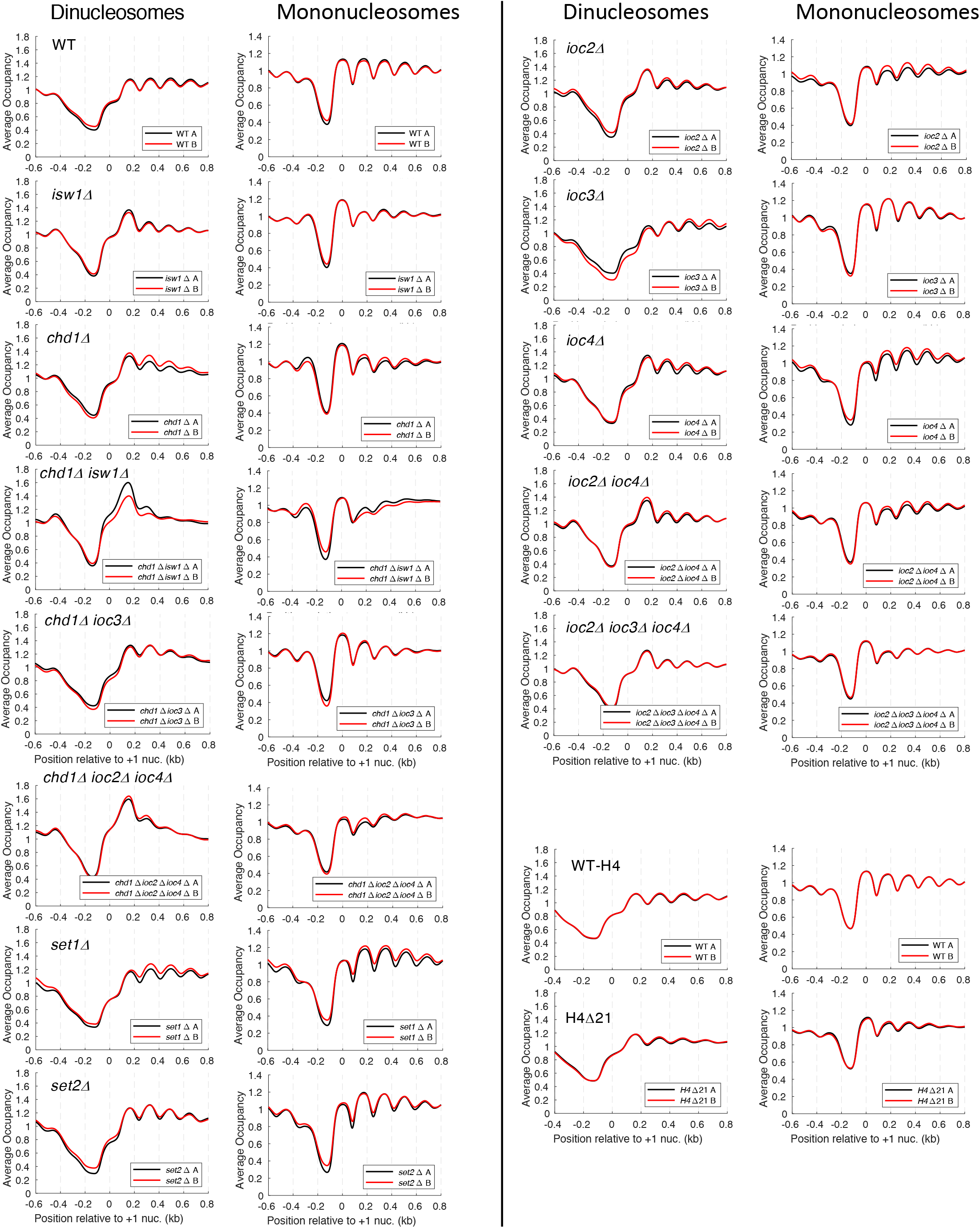
Average mononucleosome and dinucleosome occupancy (coverage) plots for all genes. Comparison of biological replicate experiments.. All 5770 yeast genes were aligned on the midpoint of their average +1 nucleosome position. Occupancy was normalised to the global average (set at 1) for dinucleosomes (250-350 bp; left panels) or mononucleosomes (120-180 bp; right panels). Note different y-axis scales are used in the dinucleosome and mononucleosome plots. Replicate A (black line); replicate B (red line).

**Supplementary Fig. S4.**
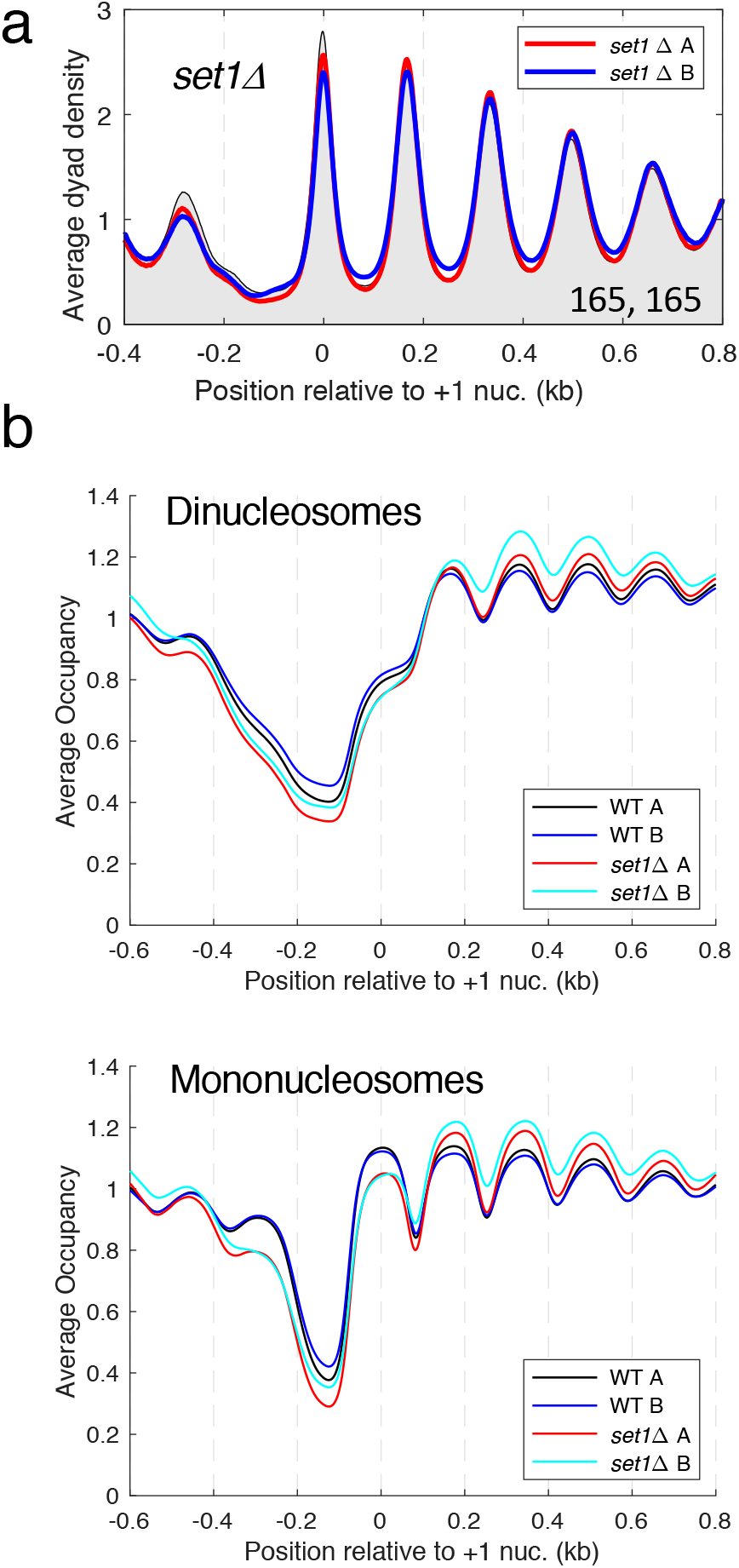
Set1 has little effect on global chromatin organisation. (**a**) Average nucleosome dyad density plot for all genes in *set1Δ* cells. Wild type replicate A is shown as a black line with grey fill. Two biological replicate experiments are shown: A (red line); B (blue line). The average spacings (bp) for replicates A and B are shown (bottom right). (**b**) Occupancy plots for dinucleosomes and mononucleosomes in *set1Δ* and wild type (WT) cells (see legend to Fig. 3).

**Supplementary Table S1.**
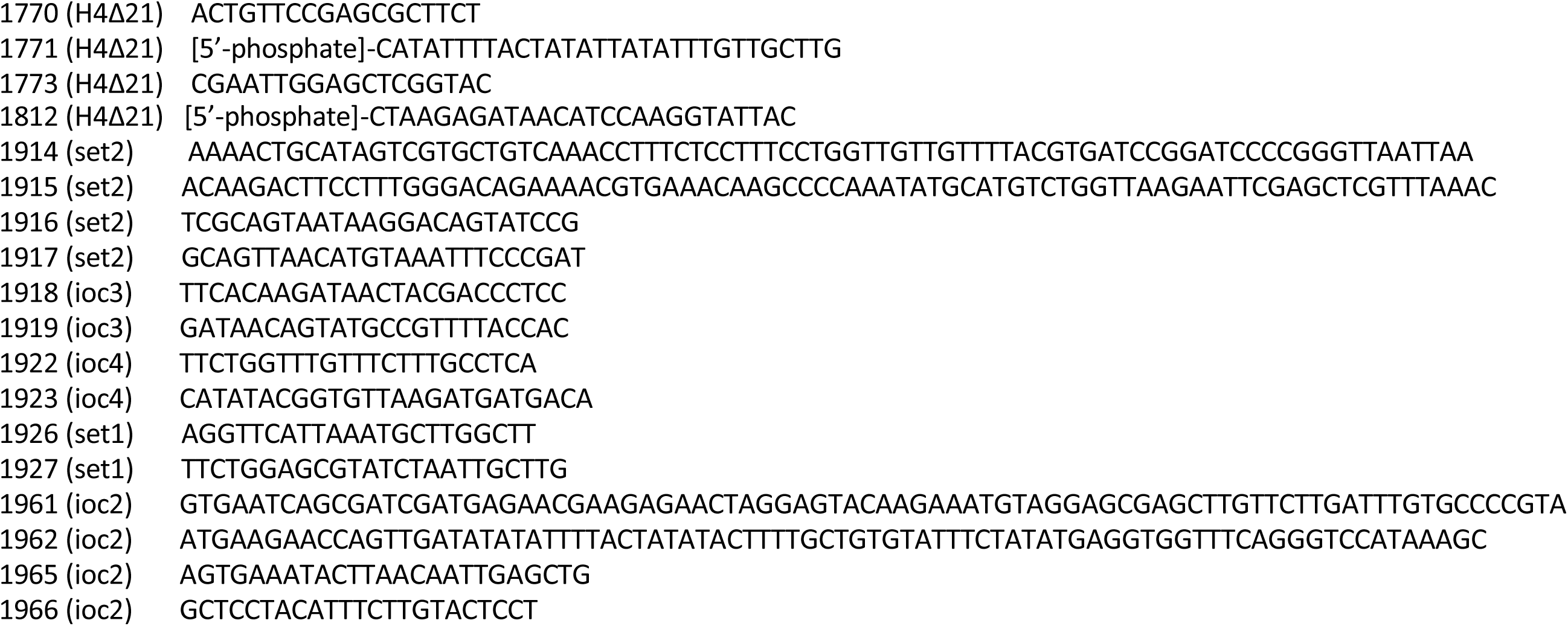
Primers used in this study.

**Supplementary Table S2.** Yeast strains used in this study. All are W303 RAD5 background (Zhao et al. 1998).

| **Strain** | **Genotype** | **Source** |
| --- | --- | --- |
| YDC111 | *MATa ade2-1 can1-100 leu2-3,112 trp1-1 ura3-1* | Kim 2006 |
| YTT186 | *MATa ade2-1 can1-100 his3-11,15 leu2-3,112 trp1-1 ura3-1 isw1Δ::ADE2* | Tsukiyama 1999 |
| YTT196 | *MATα ade2-1 can1-100 his3-11,15 leu2-3,112 trp1-1 ura3-1 isw2Δ::LEU2* | Tsukiyama 1999 |
| YTT645 | *MATa ade2-1 can1-100 his3-11,15 leu2-3,112 trp1-1 ura3-1 ioc3Δ::KanMX* | Tsukiyama Lab (unpub.) |
| YTT827 | *MATa ade2-1 can1-100 his3-11,15 leu2-3,112 trp1-1 ura3-1 ioc4Δ::HPH1* | Tsukiyama Lab (unpub.) |
| YTT1986 | *MATα ade2-1 can1-100 his3-11,15 leu2-3,112 trp1-1 ura3-1 set1Δ::NAT1* | Tsukiyama Lab (unpub.) |
| YJO482 | *MATa ade2-1 can1-100 his3-11,15 leu2-3,112 trp1-1 ura3-1 chd1Δ::HIS3MX6* | Ocampo 2016 |
| YJO484 | *MATa ade2-1 can1-100 his3-11,15 leu2-3,112 trp1-1 ura3-1 isw1Δ::ADE2 chd1Δ::HIS3MX6* | Ocampo 2016 |
| YJO486 | *MATa ade2-1 can1-100 his3-11,15 leu2-3,112 trp1-1 ura3-1 isw2Δ::LEU2 chd1Δ::HIS3MX6* | Ocampo 2016 |
| YJO487 | *MATα ade2-1 can1-100 his3-11,15 leu2-3,112 trp1-1 ura3-1 isw2Δ::LEU2 chd1Δ::HIS3MX6* | This study |
| YJO505 | *MATa/α ade2-1/ade2-1 can1-100/can1-100 his3-11,15/his3-11,15 leu2-3,112/leu2-3,112 trp1-1/trp1-1 ura3-1/ura3-1 ISW1/isw1Δ::ADE2 ISW2/isw2Δ::LEU2 RSC8/GALp-RSC8::KanMX CHD1/chd1Δ::HIS3MX6* | Ocampo 2016 |
| YPE600 | *MATa/α ade2-1/ade2-1 can1-100/can1-100 his3-11,15/HIS3 leu2-3,112/leu2-3,112 trp1-1/trp1-1 ura3-1/ura3-1 isw2Δ::LEU2* | This study |
| YPE606 | *MATa ade2-1 can1-100 leu2-3,112 trp1-1 ura3-1 ioc3Δ::KanMX* | This study |
| YPE607 | *MATα ade2-1 can1-100 leu2-3,112 trp1-1 ura3-1 ioc3Δ::KanMX* | This study |
| YPE608 | *MATa ade2-1 can1-100 leu2-3,112 trp1-1 ura3-1 ioc4Δ::HPH1* | This study |
| YPE636 | *MATa ade2-1 can1-100 leu2-3,112 trp1-1 ura3-1 ioc2Δ::URA3* | This study |
| YPE654 | *MATa ade2-1 can1-100 leu2-3,112 trp1-1 ura3-1 ioc2Δ::URA3 ioc3Δ::KanMX* | This study |
| YPE655 | *MATa ade2-1 can1-100 leu2-3,112 trp1-1 ura3-1 ioc2Δ::URA3 ioc4Δ::HPH1* | This study |
| YPE657 | *MATa ade2-1 can1-100 leu2-3,112 trp1-1 ura3-1 ioc2Δ::URA3 ioc3Δ::KanMX ioc4Δ::HPH1* | This study |
| YPE712 | *MATa ade2-1 can1-100 his3-11,15 leu2-3,112 trp1-1 ura3-1 chd1Δ::HIS3MX6 ioc3Δ::KanMX6* | This study |
| YPE715 | *MATa ade2-1 can1-100 his3-11,15 leu2-3,112 trp1-1 ura3-1 chd1Δ::HIS3MX6 ioc2Δ::URA3 ioc4Δ::HPH1* | This study |
| YPE602 | *MATa ade2-1 can1-100 leu2-3,112 trp1-1 ura3-1 set1Δ::NAT1* | This study |
| YPE604 | *MATa ade2-1 can1-100 leu2-3,112 trp1-1 ura3-1 set2Δ::TRP1* | This study |
| ROY1281 | *MATα lys2 trp1 his3 leu2 ura3 hhf1-hht1Δ::LEU2 hhf2-hht2Δ::HIS3 pCC67* | Dhillon 2000 |
| YDC507 | *MATα lys2 trp1 his3 leu2 ura3 hhf1-hht1Δ::LEU2 hhf2-hht2Δ::HIS3 p730* | This study |
| YDC101 | *MATα lys2 trp1 his3 leu2 ura3 hhf1-hht1Δ::LEU2 hhf2-hht2Δ::HIS3 p368* | This study |

